# Diacylglycerol kinase ζ deficiency triggers early signs of aplastic anemia in mice

**DOI:** 10.1101/2020.06.05.136390

**Authors:** M Martín-Salgado, E Andrada, R Liébana, M Mercedes López-Santalla, I Merida

**Affiliations:** Dept of Immunology and Oncology, National Center of Biotechnology, Madrid, Spain; Division of Hematopoietic Innovative Therapies, Centro de Investigaciones Energéticas, Medioambientales y Tecnológicas (CIEMAT), Centro de Investigación Biomédica en Red de Enfermedades Raras (CIBER-ER), Madrid, Spain; Advanced Therapy Unit, Instituto de Investigación Sanitaria Fundación Jiménez Díaz (IIS-FJD/UAM), Madrid; Spain

**Author notes:** Equal contribution (M Martín-Salgado, E Andrada). Correspondent Author: Isabel Merida. Dept of Immunology and Oncology, National Center of Biotechnology, Darwin 3, 28049 Madrid, Spain.

**Keywords:** Bone marrow, T helper 1 T cells, cytotoxic T cells, anemia, autoimmunity, diacylglycerol

## Abstract

Acquired aplastic anemia (AA) is a rare blood disorder that results from immune-mediated destruction of bone marrow (BM) progenitor cells. Improved understanding of the mechanisms that favor T cell attack in BM could help to improve early diagnosis and disease treatment. Diacylglycerol kinase ζ (DGKζ) limits T cell responses through phosphorylation of diacylglycerol into phosphatidic acid. This reaction attenuates diacylglycerol-dependent activation of the Ras/ERK/CD69 and PKCθ/NFκB pathways in response to antigen. Here we show that, in contrast to the lack of basal activation observed in peripheral lymphoid organs, DGKζ^-/-^ mice showed increased numbers of activated T cells in BM, together with a significant increase in IFNγ as well as perforin and granzyme B and C levels. The enhanced presence of T cells in DGKζ^-/-^ mouse BM correlates with reduced BM cellularity, impaired hematopoiesis, and lower frequency of circulating red cells, granulocytes, and platelets. Our studies coincide with the recent characterization of lower DGKζ expression in T cells isolated from the BM of patients with acquired AA, and suggest that limited DGKζ expression and/or functions predispose to T cell-mediated BM destruction. This study identifies the BM as a niche particularly sensitive to DGKζ deficiency and indicates that this mouse model could be of interest for studying the mechanism that contributes to AA development.

**Key points:**

- DGKζ-deficiency in mice results in larger numbers of CD69-positive T cells in bone marrow, with enhanced expression of IFNγ and lytic enzymes.
- DGKζ loss recapitulates many clinical aspects of human aplastic anemia, identifying a critical hub for immune system-dependent bone marrow failure.

**Visual abstract:** 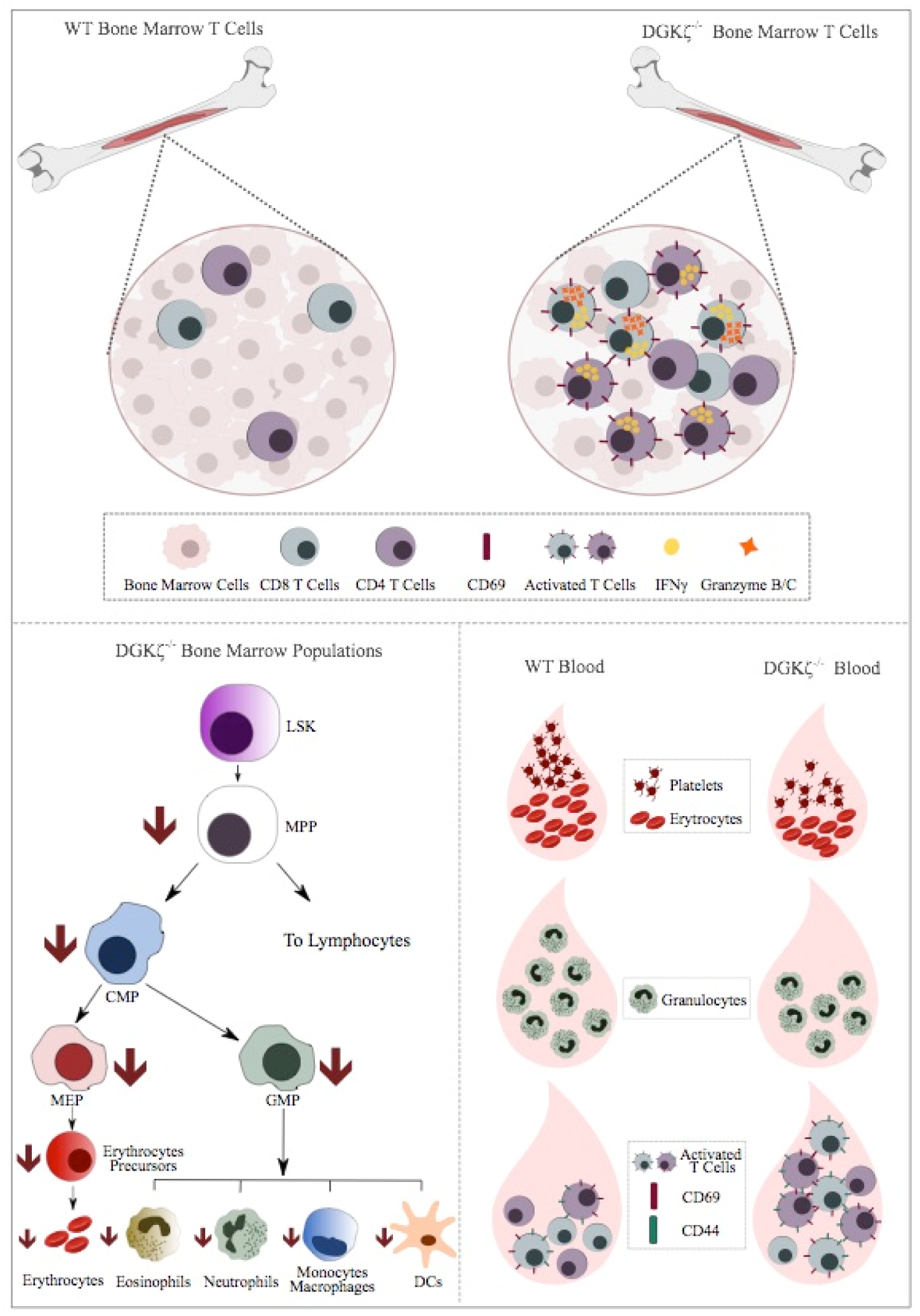

## Introduction

Aplastic anemia (AA), the prototypical bone marrow (BM) failure syndrome, is characterized by peripheral pancytopenia and hypoplastic BM.^1^ Most AA cases are acquired, idiopathic, and result from aberrant activation in the BM of T helper type-1 CD4^+^ (Th1) and cytotoxic CD8^+^ T cells (CTL).^2^ The first choice in AA treatment is allogeneic stem cell transplant using BM cells, or immunosuppressive therapies when transplant is not possible. There is an intensive search for recurrent cytogenetic abnormalities and/or somatic mutations that identify? predisposition to this disease. The identification of biological markers that indicate BM attack could assist in the selection of candidates for transplant, prognosis, and/or the predictive response to immunosuppressive therapies.

Diacylglycerol kinases (DGK) are a family of kinases that phosphorylate diacylglycerol (DAG) to phosphatidic acid (PA).^3^ DGKζ is abundantly expressed in healthy naïve peripheral T cells, where regulates DAG-dependent activation of Ras/ERK and PKCθ/NFκB signaling pathways.^4^ DGKζ downmodulation in response to antigenic challenge facilitates the correct activation of T cell helper and cytotoxic functions. Germline-DGKζ-deficient mice (DGKζ^-/-^) have hyperproliferative T lymphocytes, which are resistant to anergy induction^5^ and show enhanced elimination of xenotransplanted tumors.^6-8^

No human studies to date have described autoimmune-related DGKζ alterations. DGKζ^-/-^ mice show no overt autoimmune symptoms, which has been attributed to enhanced development and function of regulatory T cells.^9^ Upregulation of the microRNA miR-34a during T cell activation facilitates DGKζ suppression,^10^ whereas Egr2-dependent transcription triggers DGKζ induction in tumor-induced T cell anergy.^11^ Studies in AA patients have shown that mirR34a abundance in T cells is a marker of AA severity, and that higher mirR34a expression correlates with DGKζ reduction.^12^ The lesser DGKζ abundance in AA patient BM T lymphocytes suggests a causal relationship between DGKζ deficit, enhanced T cell functions, and BM destruction. This observation in AA patients prompted us to investigate the direct consequences of DGKζ genetic deletion in BM homeostasis. Our studies in DGKζ^-/-^ mice indicate increased presence of activated T cells in the BM, enhanced erythrocyte destruction, diminished hematopoietic precursor, and peripheral cytopenia. All these abnormalities, characteristic of BM failure, mirrored many of the clinical symptoms described at early stages of AA. Our studies identify an autoimmune disorder as the result of DGKζ deficiency and show the utility of DGKζ^-/-^ mice as an experimental model for the study of AA onset and development.

## Methods

### Mice

DGKζ^-/-^ and DGKα^-/-^ C57BL/6 mice were kindly donated by Drs. G Koreztky (Weill Cornell Medicine, New York, NY) and XP Zhong (Duke University Medical Center, Durham, NC). Except when indicated (4-5 weeks), animals were used between 6-12 months of age. No sex criteria were established for experiments. All procedures were conducted in accordance with Spanish and European directives and a protocol approved by the CNB/CSIC Ethic Committee for Animal Experimentation (RD53/2013).

### BM, lymph nodes, spleen and blood analysis

BM cells were extracted in phosphate-buffered saline (PBS), counted on a Countess II (ThermoFisher), and analyzed by flow cytometry or processed for RNA analysis. Lymph nodes and spleen were removed, disaggregated, and homogenized in 40 μm cell strainers (Becton Dickinson), then stained and analyzed by flow cytometry. Blood was collected by heart puncture in EDTA tubes and analyzed on an Abacus Junior Vet blood analyzer 24 h after extraction, or by flow cytometry.

### Histology

Sternum and femur bones were decalcified in 10% EDTA for 48 h or 21 days respectively, fixed in 4% paraformaldehyde, and paraffin-embedded. Tissue sections (5 μm) were stained with hematoxylin and eosin (H&E). BM cell density was analyzed using ImageJ software.

### Immunofluorescence

BM cells were lysed with RBC lysis buffer (eBioscience) to remove erythroid cells, fixed in 4% paraformaldehyde, permeabilized with 0.5% Triton X-100 (Sigma Aldrich), blocked with 10% bovine serum albumin (Sigma Aldrich), 0.05% Triton X-100 in PBS (30 min, RT), and incubated with antibodies to IFNγ (Abcam, ab9657) and CD3ε (BD Pharmingen, 550275) (ON, 4°C) in a humidified chamber. Antigen was detected using goat anti-rabbit Cy3 (Jackson) and goat anti-Armenian hamster Alexa488 (Abcam, ab173003) (60 min, RT). Images were acquired on an Olympus FV1000 confocal laser scanning microscope, and analyzed with Fiji software.

### Flow cytometry and antibodies

Antibodies used were anti-Ter119-PE, CD8-eFluor450, CD69-PeCy7, Gr1-PeCy7, Ly6G-PE (eBioscience), CD44-APC, (Coulter), CD4-PECy5, CD16/32-FITC, CD34-PE, cKit-APC, Ly6C-FITC, CD11b-PerCP5.5, Ter119-PeCy7, CD11b-PeCy7, CD4-PeCy7, CD8-PeCy7, B220-PeCy7 (BioLegend), CD69-FITC, CD62L-FITC, Sca1-biotin, NK1.1-APC, CD11c-biotin (Pharmingen), annexinV-FITC (Southern), annexinV-APC (IMMUNOSTEP) and DAPI-eFluor450. For S1P1 staining, femurs were flushed with fix solution containing 0.1% paraformaldehyde, 0.5% BSA, and 2 mM EDTA (Gibco) in PBS as reported.^13^ Cell suspensions were then lysed with RBC lysis buffer (eBioscience), incubated with rat anti-mouse S1P1-APC (R&D Systems) for 1 h on ice and washed once. Samples were then incubated (30 min, 4°C) with antibody cocktail to CD4-PercP5.5 (BioLegend), CD8-eFluor450 (eBioscience), CD44-FITC (Beckman) and CD69-PE (Pharmingen). Samples were collected on a LSRII (BD Biosciences) and analyzed with FlowJo software (FlowJo LLC, Ashland, OR).

### Induction of bone marrow failure (BMF)

F1 progeny was generated by crossing C57BL/6 and BALB/c mice. Prior to experiments F1 mice were irradiated (3 Gy,^137^Cs source); after 5 h, BMF was induced with 5 × 10^7^ splenocytes (intraperitoneal injection) from age- and gender-matched C57BL/6 wild type or DGKζ^-/-^ mice, as described.^14^ For survival studies, mice were humanely euthanized when weight loss was less than 80% of original weight.

### cDNA preparation and real time PCR

Total RNA was reverse transcribed using the High Capacity cDNA Archive Kit (PN4368813; Applied Biosystems). Real-time PCR reactions were performed in triplicate with HOT FIREPol EvaGreen qPCR Mix Plus (ROX) (Solis BioDyne) in an Applied Biosystems ABI PRISM 7900HT machine with SDS v2.4 software, using a standard protocol. Results were analyzed by the comparative Ct method (ΔΔCt). Expression was normalized using the β-actin housekeeping gene for each sample. Primers used were:IL-6:5’-GCTACCAAACTGGATATAATCAGGA-3’ and 3’-AAGACCTCATGTATCGATG GACC-5’; TNF-α: 5’-CCCTCACACTCAGATCATCTTCT-3’ and 3’-GACATCGGGTGCA GCATCG-5’. TGF-β: 5’-ACCATGCCAACTTCTGTCTG-3’; 3’-AGATGTTGGTTGTGTT GGGC-5’. Primers for IFNγ and perforin,^15^ for granzymes A, B and C,^16^ for IL-10 and IL-2,^17^ for KLF-1, cMpl and p57,^18^ and for β-actin^19^ as reported.

### Statistical analysis

Data were analyzed using GraphPad Prism 5. Results are represented as mean ± standard error of the mean (SEM). Normality was analyzed by the Kolmogorov-Smirnov test. An unpaired two-tailed t-test with 95% confidence intervals was used for data with normal distribution and equal variances, and an unpaired t-test with Welch’s correction for data sets with different variances. The Mann-Whitney U test was used for data with non-normal distribution. For more than two groups with normal distribution, ANOVA with Bonferroni correction was performed. Kruskal-Wallis test with Dunn’s post hoc was applied when two groups with non-normal distribution. Survival statistics were performed with the Gehan-Breslow-Wilcoxon test. Differences were considered non-significant (ns) when *p* >0.05, significant (*) when *p* <0.05, very significant (**) when *p* <0.01 and extremely significant (***) when *p* <0.001.

## Results

DGKζ deficiency in mice causes no gross defects in T cell development and leads to a slight decrease in mature T cells in spleen, with no signs of activation in basal conditions.^5^ The BM from young (4- to 5-week old) DGKζ^-/-^ mice showed similar total numbers of CD4^+^ and CD8^+^T cells compared to WT mice (Figure 1A). CD69 is an early activation marker that identifies recent antigen-stimulated T lymphocytes.^20^ Both T cell pools in the BM of DGKζ^-/-^ mice showed significantly augmented C69 populations with heightened CD69 expression on a per cell basis compared to WT animals (Figure 1B). A greater number of CD69^+^ T lymphocytes win DGKζ^-/-^ mouse BM correlates with observations in AA patients, whose BM CD69^+^ T cell numbers are greater than those of healthy controls.^12^

**Figure 1.**
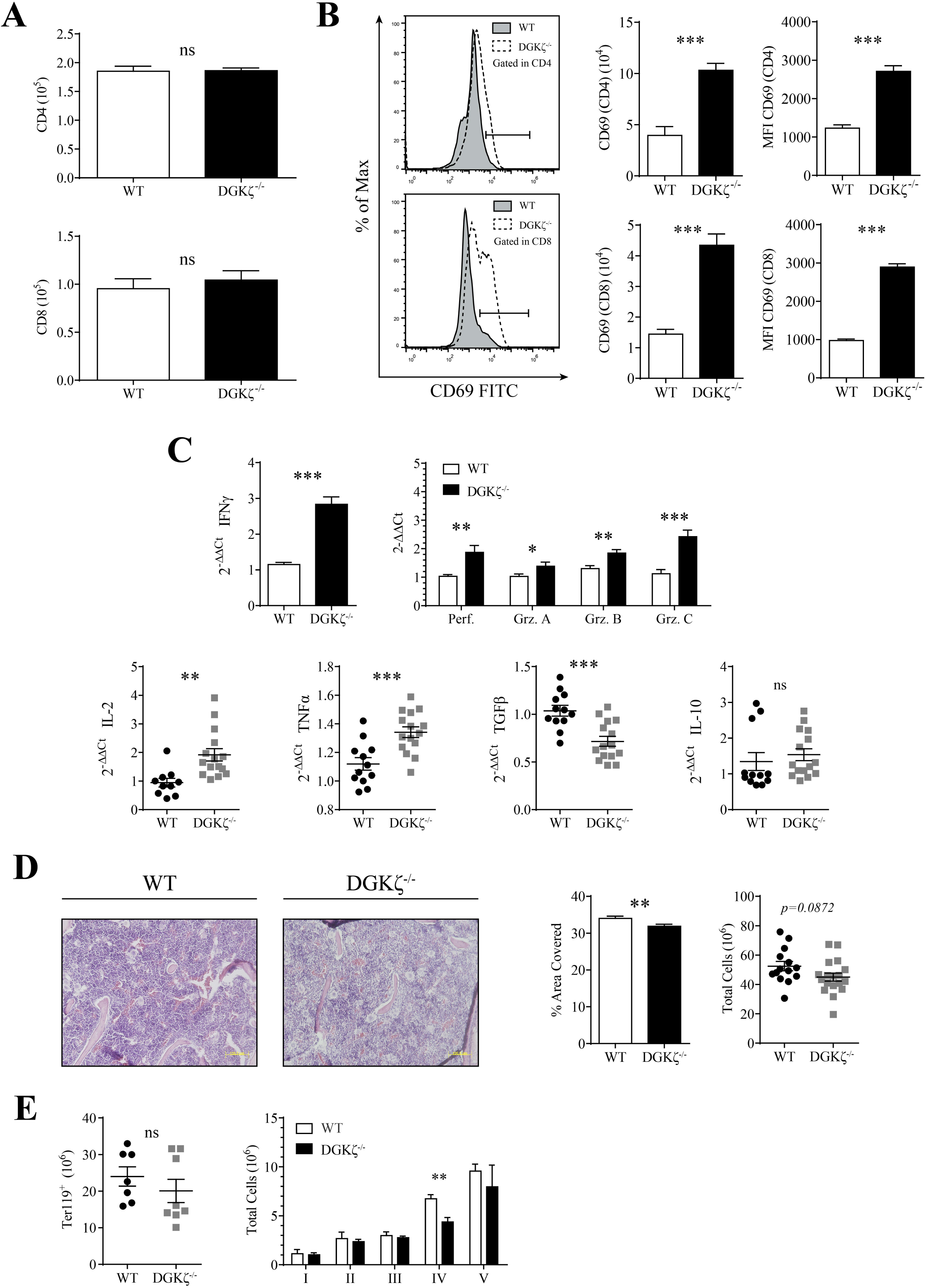
Analysis of BM T cells from DGKζ deficient mice. BM cells from WT and DGKζ^-/-^ mice were flushed from tibias and femurs and processed for flow cytometry analysis. **(A)** Quantification of CD4^+^ and CD8^+^ T cells counts in the BM of 4- to 5-week old mice (gated in total cells). **(B)** Left, representative histograms of CD69 expression gated on CD4^+^ or CD8^+^ T cell populations. Middle, total counts of activated T cells CD69^+^ CD4^+^ and CD69^+^ CD8^+^ cells. Right, mean fluorescence intensity (MFI) of CD69 in CD4^+^ and CD8^+^ populations. **(C)** mRNA expression was analyzed from total BM cells by RT-qPCR. Top, expression of IFNγ, perforin and granzymes (Grz.). Bottom, mRNA levels of IL-2, TNFα, TGFβ and IL-10. **(D)** Left, femur sections were fixed, and H&E stained. Images show central area of each section (200x). MiddleCenter, quantification of the percentage of area covered by cells. Right, quantification of total BM cells by TrypanBlue exclusion. **(E)** Left, quantification of total erythroid cells in WT and DGKζ^-/-^ mice by Ter119 expression. Right, analysis of erythroid differentiation. A. Mean ± SEM. n=6/genotype. CD4^+^. Unpaired t-test. CD8^+^. Mann-Whitney test. B. Mean ± SEM. n=6/genotype. Unpaired t-test. C. Mean ± SEM. WT n=4; DGKζ^-/-^ n=5. IFNγ, perforin and Grz. C. Unpaired t-test with Welch’s correction. Grz. A, Grz. B, IL-2, TNFα and TGFβ. Unpaired t-test. IL-10. Mann-Whitney test. Data were acquired in two independent experiments. D. Mean ± SEM. Left and center, four sections/mouse; WT n=4, DGKζ^-/-^ n=5. Unpaired t-test. For total cellularity, WT n=14, DGKζ^-/-^ n=18. Unpaired t-test. E. Mean ± SEM. WT n=7, DGKζ^-/-^ n=8. E. Mann-Whitney test. Data were acquired in two independent experiments.

Expanded numbers of IFNγ-expressing Th1 T cells is an indicator of AA,^2^ and low IFNγ^+^ T cell frequency correlates with responsiveness to immunosuppressive therapy.^21^ mRNA expression analysis showed a significant increase in IFNγ abundance in DGKζ^-/-^ BM compared to WT mice (Figure 1C). Additional analysis showed higher expression of perforin and granzymes A, B and C as well as IL-2 and TNFα, with reduction in TGFβ expression (Figure 1C). Greater IFN-γ and TNFα expression correlates well with the enhanced skewing toward Th1 phenotypes that results from DGKζ deficiency,^22^ and resembles the increased levels of circulating IFNγ in AA patients.^23^ Augmented expression of granzymes B and C and perforin in DGKζ^-/-^ mouse BM denotes an increase in cytotoxic T cells (CTL) similar to that described in AA patients.^24^ Activated T lymphocytes cause BM destruction in AA;^25^ DGKζ^-/-^ mice showed reduced BM cellularity (Figure 1D). Analysis of the TER-119^+^ erythroid lineage populations did not show differences between DGKζ^-/-^ and WT mice (Figure 1E). Nonetheless, combined analysis of CD44 expression as a function of cell size in the TER-119^+^ population^26^ showed lower cell numbers at stage IV in DGKζ^-/-^ mice (Figure 1E).

The BM failure in AA is attributed in great measure to the abnormal presence of inflammatory cytokines that promote HSC differentiation, preventing their continuous self-renewal.^27^ HSC activity in the BM is found in a small subset of the lineage marker (Lin)^-^ Sca1^+^c-Kit^+^ (LSK) population that gives rise to multipotent progenitors (MMP).^28^ Immunophenotype analysis of HSC by flow cytometry showed no differences in the LSK pool in DGKζ^-/-^ mice, with a significant decrease in MMP compared to WT mice (Figure 2A). Further analysis revealed a reduction in the common myeloid progenitors (CMP) as well as megakaryocyte/erythrocyte progenitors (MEP) and granulocyte/monocyte progenitors (GMP) (Figure 2B). Analysis of some key hematopoietic factors showed severely reduced expression of Kruppel-like factor 1 (KLF-1), essential for erythroid lineage commitment and maturation in mice and humans^29^ (Figure 2C). DGKζ^-/-^ mice also showed lower expression of the thrombopoietin (THPO) receptor c-Mpl (Figure 2C). THPO binding to its receptor sustains adult quiescent HSC populations by regulating cyclin-dependent kinase inhibitors including p57^kip2^.^30,31^ Reduced p57^kip2^ expression pointed to impaired HSC quiescence in DGKζ^-/-^ mice (Figure 2C).

**Figure 2.**
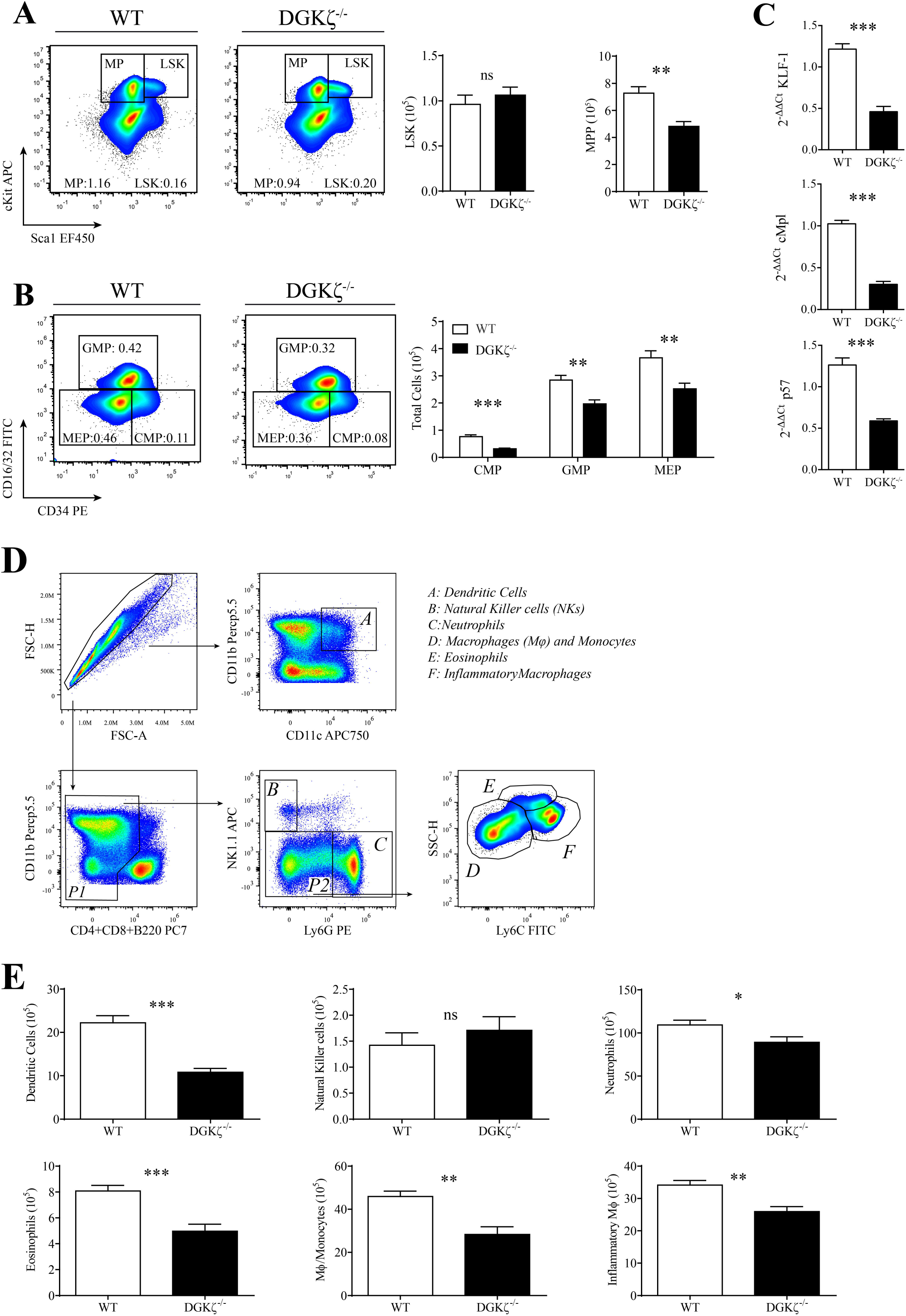
DGKζ^-/-^ mice show defects in HSCs and myeloid development in the BM. **(A)** Left, representative flow cytometry dot plots of cKit and Sca1. Right, quantification of total of LSK cells (Lin^-^ cKit^+^ Sca1^+^) (gated on Lin^-^ BM cells), and multipotent progenitors (MP) (Lin^-^ cKit^+^ Sca1^-^) (gated on Lin^-^ cells). **(B)** HSCs composition was gated on Lin^-^ BM cells by flow cytometry. Left, representative dot plots of CD16/32 and CD34. Right, quantification of common myeloid progenitors (CMP) (Lin^−^ cKit^+^ Sca1^−^ CD34^+^ CD16/32^int^), granulocyte/monocyte progenitors (GMP) (Lin^−^ cKit^+^ Sca1^−^ CD34^+^ CD16/32^hi^), and megakaryocyte/erythrocyte progenitors (MEP) (Lin^−^cKit^+^Sca1^−^CD34^−^CD16/32^lo^) (gated on MP). **(C).** mRNA was purified from total BM cells and KLF1, cMpl and p57 expression was analyzed by RT-qPCR. **(D)** Flow cytometry dot plots of gating strategy followed for myeloid characterization. **(E)** Quantification of total dendritic cells (CD11b^+^ CD11c^+^), natural killer cells (NK1.1^+^ Ly6G^-^) (gated on CD11b^low/high^ CD4^-^ CD8^-^ B220^-^), neutrophils (NK1.1^-^ Ly6G^+^) (gated on CD11b^low/high^ CD4^-^ CD8^-^ B220^-^), eosinophils (Ly6C^dim^) (gated on CD11b^low/high^ CD4^-^ CD8^-^ B220^-^ NK1.1^-^ Ly6G^low/dim^), macrophages and monocytes (Ly6C^low^) (gated on CD11b^low/high^ CD4^-^ CD8^-^ B220^-^ NK1.1^-^ Ly6G^low/dim^), and inflammatory macrophages (Ly6C^high^) (gated on CD11b^low/high^ CD4^-^ CD8^-^ B220^-^ NK1.1^-^ Ly6G^low/dim^). A, B. Mean ± SEM. WT n=5; DGKζ^-/-^ n=7. Unpaired t test. Data were acquired in two independent experiments. C. Mean ± SEM. WT n=4; DGKζ^-/-^ n=5. 2^-ΔΔCt^ p57. Unpaired t-test with Welch’s correction. 2^-ΔΔCt^ KLF1 and cMpl. Unpaired t-test. D, E. Mean ± SEM. n=7/genotype. Unpaired t-test. Data were acquired in two independent experiments.

We analyzed mature myeloid populations using antibodies to target cell surface proteins, in combination with forward *vs* side scatter (FSC and SSC) parameters as described.^32^ Dendritic subsets (CD11c^h^ CD11b^+^) were differentiated from other CD11b^+^ myeloid subsets and were further analyzed for NK1.1^+^ natural killer (NK) and Ly6G^+^ neutrophil subsets (Figure 2D). Eosinophils (Ly6C^Int^Ly6G^-^SSC^hi^) were differentiated from the monocyte/macrophage population that were further analyzed for the presence of Ly6C^+^ and more mature Ly6C^low/neg^ populations.^33^ DGKζ^-/-^ mouse BM showed a substantial reduction in all myeloid populations except NK cells (Figure 2E).

These results demonstrate that young DGKζ^-/-^ mice have activated T cells in BM and a concurrent reduction in precursors of erythroid and myeloid populations. Lineage tracing experiments show that HSC produce the majority of conventional myeloid cells and platelets within 8 months, approximately half of the mouse lifespan.^34^ We thus explored the evolution of BM populations in adult (6-12 months) WT and DGKζ^-/-^ mice. Total CD4^+^ T cell numbers in the BM from DGKζ^-/-^ mice were elevated compared to those in WT mice (Figure 3A). CD69 analysis confirmed increase both in the total numbers of CD69 positive and the CD69 expression in CD4 and CD8 T cell populations (Figure 3B). In addition to acting as a T cell activation marker, CD69 functionally promotes BM retention by binding to and sequestering the sphingosine-1-phosphate receptor (S1P1), necessary for BM egress.^35^ In accordance, the percentage of S1P^+^ CD4^+^ T cells was lower in DGKζ^-/-^ compared to controls. A lesser, non-significant tendency was observed in CD8^+^ T cells (Figure 3C).

**Figure 3.**
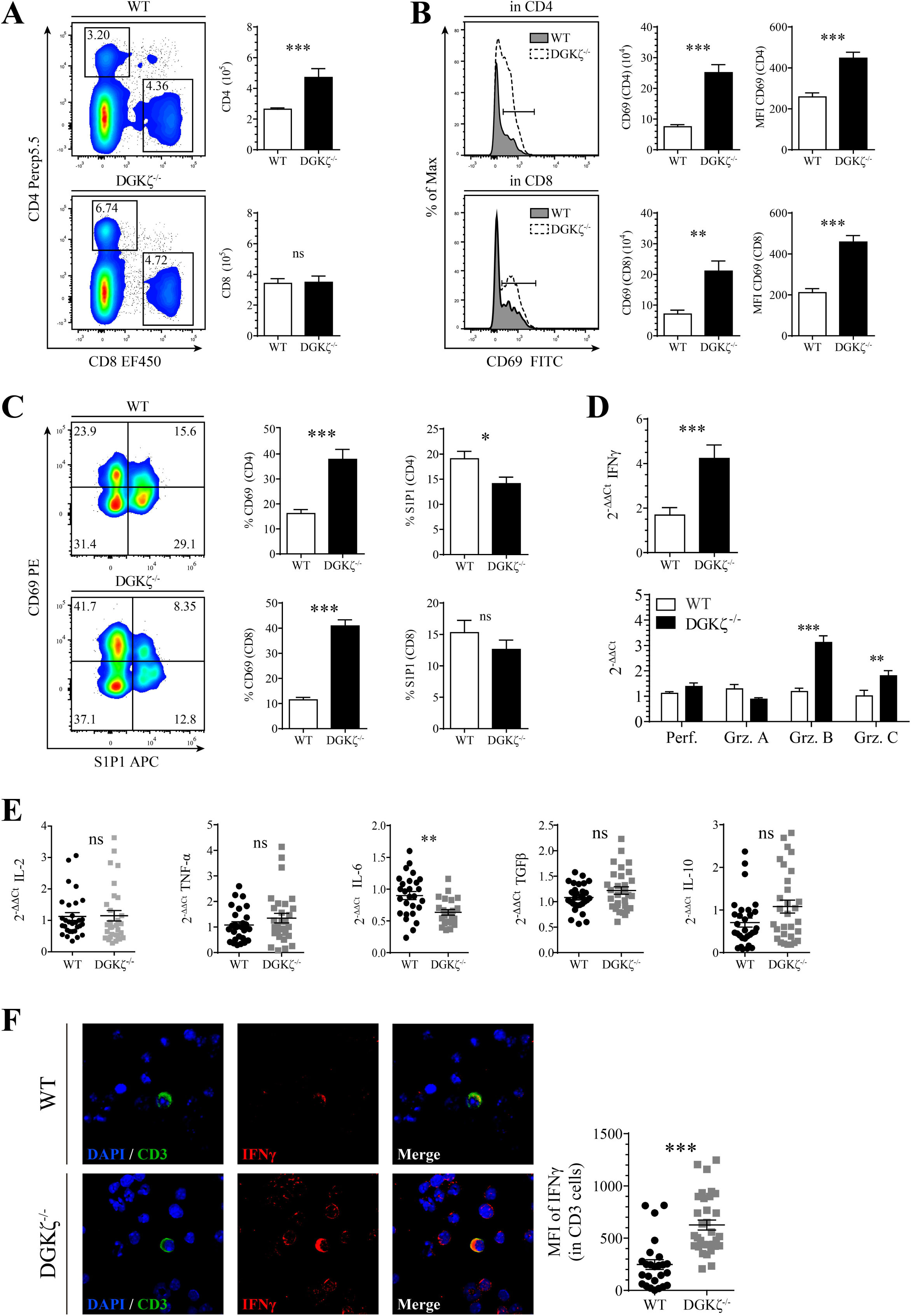
Analysis of BM T cells of mature DGKζ^-/-^ mice. **(A)** Left, flow cytometry dot plots of CD4 and CD8 T cell populations in total BM. Right, quantification of total CD4 and CD8 T cells. **(B)** Left, representative histograms of CD69 expression gated either on CD4^+^ or CD8^+^ T cell populations. Middle, total numbers of activated T cells CD69^+^CD4^+^ and CD69^+^CD8^+^ T cells. Right, MFI of CD69 gated on CD4^+^ or CD8^+^ T cells **(C)** S1P1 expression was analyzed in T cells from WT and DGKζ^-/-^ mice by flow cytometry. Left, representative flow cytometry dot plots of CD69 vs S1P1. Right, percentages of CD69^+^ and S1P1^+^ T cells (gated on CD4^+^ or CD8^+^). **(D, E)** mRNA expression of IFNγ, perforin (Perf), granzymes (Grz) and interleukin (IL) 2, 6, 10 and TNFα was analyzed by RT-qPCR from total BM lysates. **(F)** CD3ε and intracellular IFNγ were stained and analyzed by confocal microscopy in total BM cells. Bottom, quantification of MFI of IFNγ in CD3ε cells. A, B. Data shown as mean ± SEM. WT n=13; DGKζ^-/-^ n=14. A. Mann-Whitney test. B. Unpaired t-test with Welch’s correction (for CD69^+^ CD4^+^ and CD69^+^ CD8^+^), Mann-Whitney test (for MFI CD69 in CD4) and unpaired t test (for MFI CD69 in CD8). Data were acquired in three independent experiments. C. Mean ±SEM. n=15/genotype. For % CD69. Unpaired t test with Welch’s correction. For % S1P1. Unpaired t test. Data were acquired in three independent experiments. D, E. Mean ± SEM. n=10/genotype. Perf., Grz. A, B, C, IL-2 and IL-10. Mann-Whitney test. TNFα, IL-6 and TGFβ. Unpaired t-test with Welch’s correction. Data were acquired in three independent experiments. F. Mean ± SEM. Eleven images/mouse. n=3/genotype. Mann-Whitney test.

Similar to that observed in young mice, mRNA analysis indicated augmented expression of IFNγ and granzymes B and C (Figure 3D). No differences were observed for IL-2, TNFα, TGFβ and IL-10, except for lower IL-6 expression (Figure 3E). Immunofluorescence analysis confirmed increased mean fluorescence intensity (MFI) of IFNγ in CD3^+^ cells from BM of DGKζ^-/-^ compared to WT mice (Figure 3F).

Analysis of BM from mature WT and DGKζ^-/-^ showed substantial loss of cellularity (Figure 4A) that correlated at this age with a marked reduction of the erythroid TER-119^+^ population^36^ (Figure 4B). Annexin V staining suggested a higher apoptosis rate in DGKζ^-/-^ TER-119^+^erythrocytes compared to WT mice (Figure 4B). Based on CD44 vs. FSC gating for erythroid differentiation, DGKζ^-/-^ showed reduced cell numbers at all differentiation stages (Figure 4C). Annexin V staining for apoptosis detection was higher at stage I, the earliest progenitors, and at stage IV, where erythroid precursors show the greatest sensitivity to apoptotic cell death^37^ (Figure 4C). Analysis of cells from the myeloid lineage in DGKζ^-/-^ mice showed reduced numbers of neutrophils and monocytes/macrophage populations (Figure 4D).

**Figure 4.**
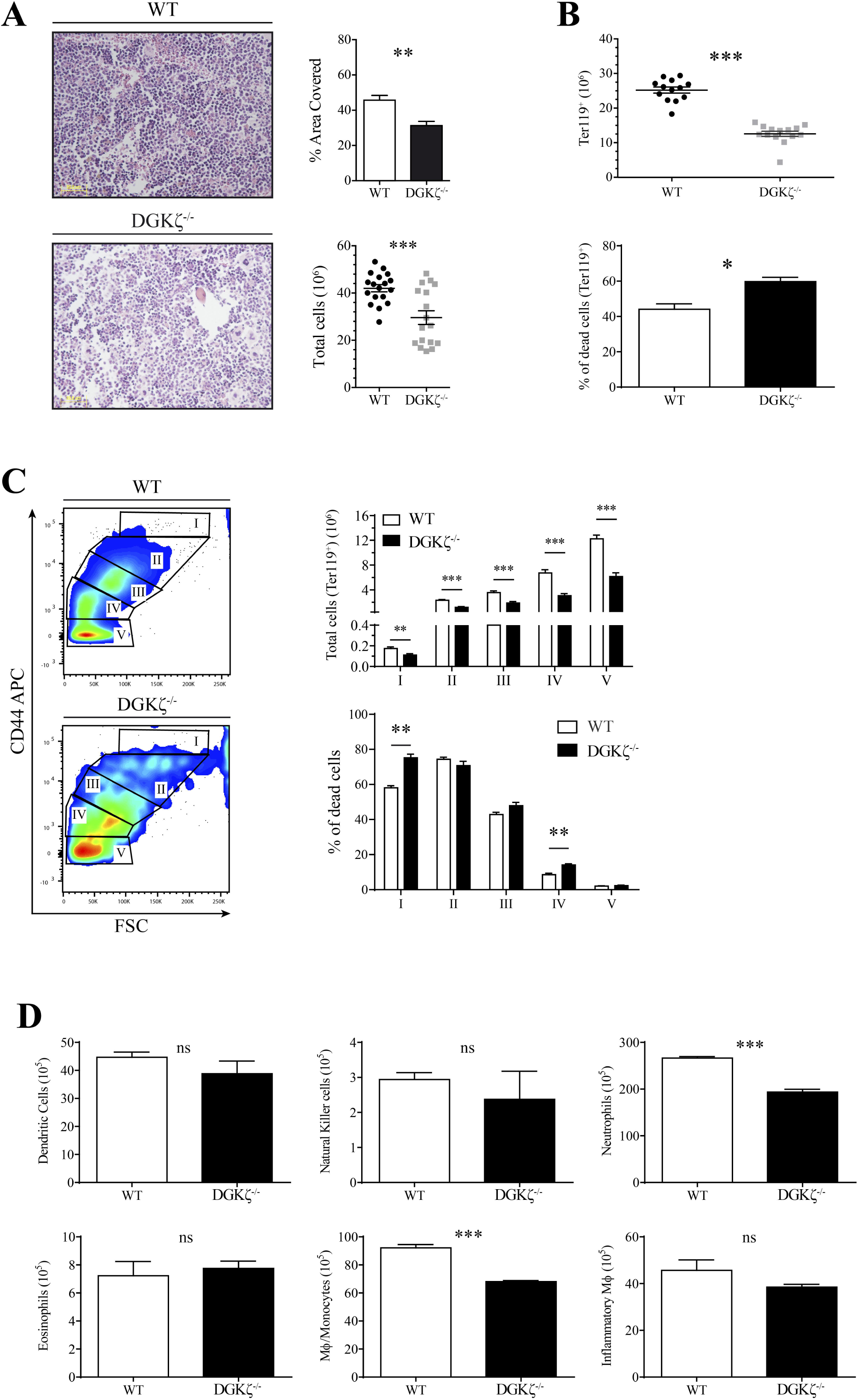
DGKζ^-/-^ mice show signs of BM aplasia and defects in the erythroid compartment. **(A)** Left, femurs from WT and DGKζ^-/-^ mice were fixed and H&E stained. Images show central area of each section (200x). Top right, quantification of the percentage of area covered by cells in BM samples. Bottom right, BM cells obtained from tibias and femurs of WT and DGKζ^-/-^ mice were quantified by TrypanBlue exclusion. **(B)** Total erythroid lineages and different erythroid populations within the BM were analyzed by flow cytometry. Top, total erythroid counts (Ter119^+^) (gated on total cells). Bottom, percentage of cell death Ter119^+^ population (% annexinV^+^ cells + % annexinV^+^ DAPI^+^ cells). **(C)** Left, representative dot plots of CD44 and FSC populations in WT and DGKζ^-/-^ mouse BM (gated on Ter119^+^ cells). Top right, quantification of erythroid populations (in Ter119^+^ cells). Bottom right, percentage of cell death (% annexinV^+^ cells + % annexinV^+^ DAPI^+^ cells) for the different erythroid populations. **(D)** Quantification of total dendritic cells (DC), natural killer cells (NK), neutrophils, macrophages (Mφ)/monocytes, eosinophils and inflammatory Mφ in the BM of WT and DGKζ^-/-^ mice following the gating strategy depicted in Figure 2. A. Data shown as mean ± SEM. Three sections/mouse; n=3/genotype. Mann-Whitney test. For total cellularity, n=14/genotype. Unpaired t-test with Welch’s correction. B. Mean ± SEM. Top, n=14/genotype. Mann-Whitney test. Bottom, n=3/genotype. Unpaired t-test. C. Mean ± SEM. Top right, n=14/genotype. Mann-Whitney test. Bottom right, n=3/genotype. Unpaired t-test. Data were acquired in three independent experiments. D. Data shown as mean ± SEM. n=3/genotype. Unpaired t-test. Data were acquired in a single experiment.

DGKα is another DGK family member expressed highly in T lymphocytes and has partially redundant functions in the control of Ras/ERK signals downstream of the TCR.^4^ At difference from the case for DGKζ, DGKα^-/-^ mice showed normal numbers of BM CD4 and CD8 T cells (Suppl Figure 1A) with no differences in CD69 expression (Suppl Figure 1B). Ter119^+^ populations were not altered (Suppl. Figure 1C), with similar cell numbers at all differentiation stages (Suppl. Figure 1D). These results strongly suggest an isoform-specific DGKζ contribution limiting T cell activation at the BM.

Immune-dependent destruction of BM precursors together with continuous differentiation eventually translates into peripheral cytopenia in AA patients. Blood analysis of DGKζ^-/-^ mice showed reduction in RBC counts, as well as decreased hematocrit and hemoglobin content compared to WT mice (Figure 5A). Similar lymphocyte counts (Figure 5B) contrasted with severe reduction in circulating granulocytes and platelets, as anticipated from BM analysis (Figure 5C). These data fully agree with the impaired generation of precursors, erythrocyte and myeloid populations, and correlate with clinical symptoms observed in AA patients.

**Figure 5.**
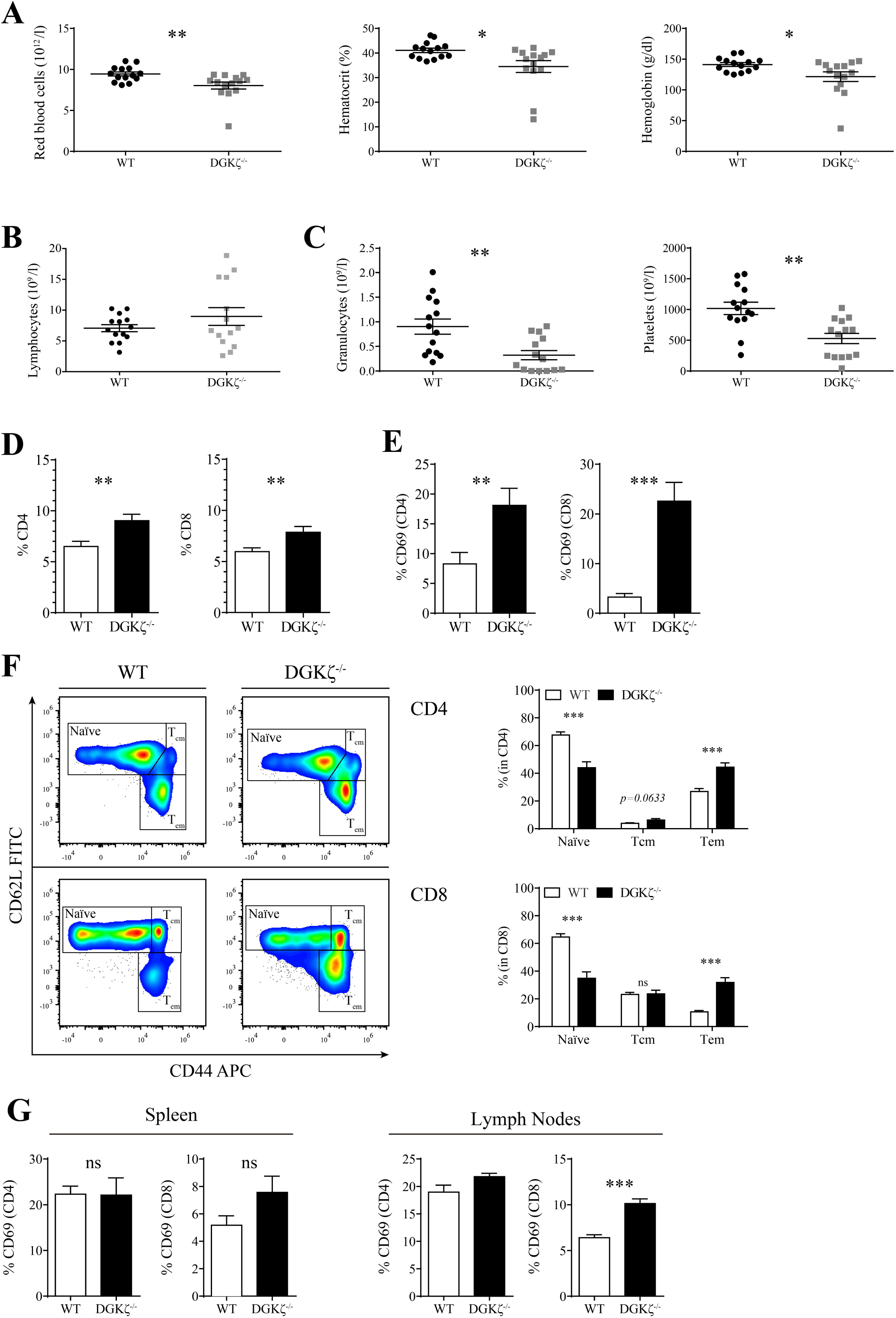
Analysis of peripheral blood of DGKζ^-/-^ mice. Blood was collected by heart puncture from WT and DGKζ^-/-^ mice and analyzed on an Abacus Junior Vet blood analyzer or processed for flow cytometry. **(A)** Left, analysis of total circulating red blood cells. Center, hematocrit analysis. Right, hemoglobin quantification. **(B)** Total circulating lymphocytes. **(C).** Left, analysis of total granulocytes. Right, analysis of circulating platelets. **(D)** Flow cytometry analysis of blood samples for circulating T lymphocyte characterization. Percentages of CD4^+^ and CD8^+^ T cells (gated on total cells). **(E)** Percentages of CD69^+^ (gated on CD4^+^ or CD8^+^). **(F)** Left, representative flow cytometry dot plots of CD62L and CD44 cells gated on CD4 (top) or CD8 (bottom). Right, quantification of percentages of CD62L^high^CD44^low^ (naïve), CD62L^high^CD44^high^ (central memory, T_cm_) and CD62L^low^CD44^high^ (effector memory, T_em_) gated on CD4^+^ or CD8^+^. **(G)** CD69 expression was analyzed in spleen and lymph nodes. A-C. Mean ± SEM. n=14/genotype. A. Left, Unpaired t-test. Center, Mann-Whitney test. Right, Unpaired t-test with Welch’s correction. B. Unpaired t-test with Welch’s correction. C. Left, Unpaired t-test. Right, Mann-Whitney test. Data were acquired in three independent experiments. D-F. Mean ± SEM. n=13/genotype. D. Percentage of CD4. Mann-Whitney test. Percentage of CD8. Unpaired t-test. E. Percentage of CD69 (CD4). Unpaired t-test. Percentage of CD69 (CD8). Mann-Whitney test. Percentage of CD62L^high^CD44^low^ (naïve), CD62L^high^CD44^high^ (central memory, T_cm_) and CD62L^low^CD44^high^ (effector memory, T_em_) in CD4. Mann-Whitney test. In CD8. Mann-Whitney test. Data were acquired in four independent experiments. G. Mean ± SEM. n=3/genotype. Unpaired t-test.

**Figure 6.**
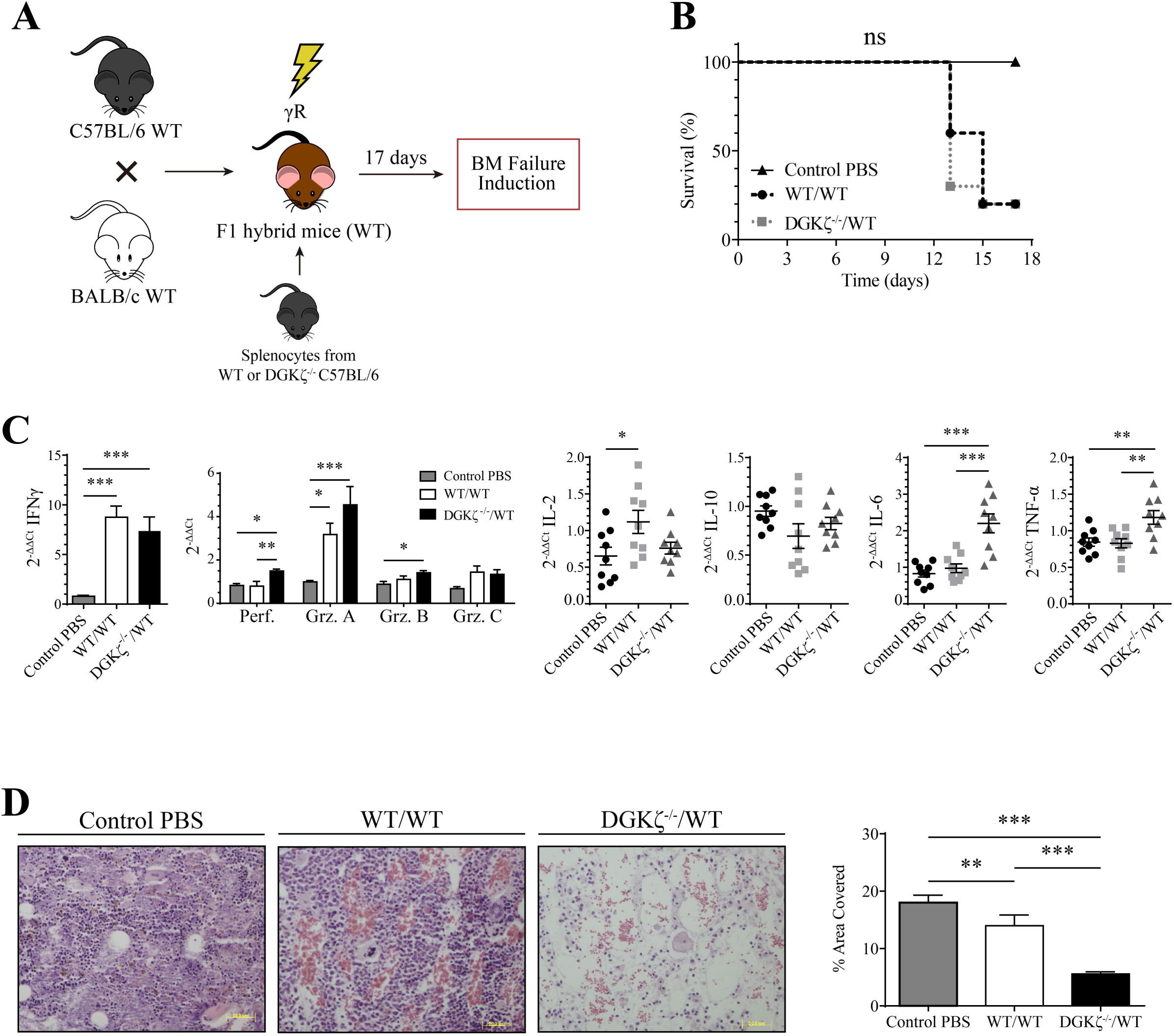
DGKζ deficiency enhances immune-dependent BM destruction. (**A**) Scheme for the generation of a mouse model of severe aplastic anemia: C57BL/6 (Hb/b) and BALB/c (Hd/d) mice were crossed and F1 progeny (Hb/d) mice were sublethally irradiated (γR) and infused with 5×10^7^ WT or DGKζ^-/-^ C57BL/6 splenocytes by i.p. injection. (**B**)Kaplan–Meier survival curve (day +17) estimates percentage of living mice after aplastic anemia induction with WT or DGKζ^-/-^ C57BL/6 splenocytes. **(C)** mRNA expression of Grz A, B, C, perforin, IFNγ, IL-2, TNFα, IL-10 and IL-6 was analyzed by RT-qPCR on day +7 after bone marrow failure induction. **(D)** Left, sternums from F1 hybrid mice were fixed and H&E stained on day +17 post disease induction. Images show central area of each section (200x). Right, quantification of the percentage of area covered by cells in sternum samples after aplastic anemia induction. B. N=10/genotype; control PBS n=·2. Gehan-Breslow-Wilconxon test. Data were acquired in two independent experiments. C. Mean ± SEM. n=3/genotype. One-way ANOVA with the Bonferroni post hoc test. D. Mean ± SEM. Three sections/mouse; WT/WT n=8; DGKζ^-/-^/WT n=10; control PBS n=2. Kruskal-Wallis test and Dunn’s post hoc. One representative image of each sample is shown. Data were acquired in two independent experiments.

As human studies show that the most activated T cells enter the circulation,^38^ we analyzed the activation profile of WT and DGKζ^-/-^ peripheral blood cells (PBL). DGKζ^-/-^ mice showed a higher percentage of circulating CD4 and CD8 T cells (Figure 5D), with large increase in CD69^+^CD4^+^ and CD8^+^ T cells (Figure 5E). Based on L-selectin (CD62L) and CD44 expression, we analyzed blood for CD62L^h^CD44^lo^ naive (TN), CD62^lo^CD44^h^ effector memory (TEM) and CD62^h^CD44^h^ central memory (TCM) T cells (Figure 5F); in agreement with heightened expression of the activation marker CD69, blood analysis from DGKζ^-/-^ mice showed reduced abundance of TN populations with an increased percentage of TEM (Figure 5F) and TCM CD4^+^ T compared to WT mice (Figure 5F). In agreement with other reports,^5^ CD69 expression was similar in DGKζ^-/-^ and WT splenocytes and lymph nodes (Figure 5G). The analysis of DGKζ^-/-^ PBL reflects the activation T cell phenotype observed in the BM. The activation phenotype is reminiscent of the reduced frequency of TN CD4^+^/CD8^+^ cells^39^ and the increase percentages of CD4^+^ and CD8^+^ TEM subsets ^40^ that are described in PBLs of AA patients.

All these experiments suggested direct correlation between DGKζ loss and enhanced T cell-mediated destruction of the BM. To further demonstrate a causal relationship between DGKζ deficiency in the T cell compartment and BM damage, we used a well-characterized murine model of immune-dependent BM failure.^14^ Severe AA was induced via sublethal irradiation of F1 hybrid mice, followed by adoptive transfer of WT or DGKζ^-/-^ C57BL/6 splenocytes (Figure 7A). All infused animals showed symptoms rapidly and were sacrificed between 14 and 17 days (Figure 7B). In accordance with the higher IFNγ serum concentrations detected in murine lymphocyte infusion-induced BM failure models,^41^ the BM of mice analyzed 7 days after infusion with WT splenocytes showed enhanced IFNγ expression compared to irradiated control animals (Figure 7C). Augmented granzyme A and IL-2 expression was in agreement with the massive infiltration of CD8^+^ (NK and T cells) described in this infusion model.^41^ Compared to mice infused with WT splenocytes, hosts receiving DGKζ^-/-^ cells showed higher expression of perforin and granzymes A and B indicative of strong cytotoxic function (Figure 7C). The lack of IL-2 increase observed in hosts infused with DGKζ-deficient cells, contrasted with enhanced expression of TNF-α and IL-6 (Fig 7C). In mouse models of graft-versus-host disease (GVHD), TNFα production correlates with the destructive phase of the disease.^42^ Enhanced IL-6 production is also an important factor in the severity of murine GVDH pathology.^43^ In accordance with the recognized contribution of TNFα and IL-6, the extent of BM destruction was higher in F1 mice that received DGKζ^-/-^ cells compared to those infused with WT cells (Figure 7D).

## Discussion

Studies in animal models have shown that antigen recognition in the absence of DGKζ results in stronger Ras/PKCθ activation, leading to enhanced AP-1/NFκB-regulated transcription.^44^ This promotes a CD4 Th1/ Th2 skew that provides DGKζ^-/-^ mice with enhanced anti-viral responses,^5^ granting protection against allergic asthma.^22^ DGKζ deficiency also potentiates cytotoxic programs in CD8^+^ T and NK cells, boosting antitumor functions in an antigen-dependent ^45^ and -independent manner.^6,46^ These and other studies prompted a growing interest in DGKζ as a pharmacological target with immunomodulator abilities. A central issue when considering the therapeutic potential of negative regulators of T cell function is the risk of unleashing adverse autoimmune responses. Characterization of autoimmune disease symptoms in DGKζ^-/-^ mice is critical to fully assessing the therapeutic potential of this enzyme.

Our studies identify abnormal accumulation of CD69-positive T cells in the BM of untreated DGKζ^-/-^ mice, a situation reminiscent of the enhanced CD69 expression observed in BM T lymphocytes from AA patients. The lower cellularity, impaired erythropoiesis, and deficient progenitor population numbers found in DGKζ^-/-^ mouse BM suggest immune-dependent BM failure similar to that described in AA. The observation of diminished DGKζ expression in the T cells from the BM of AA patients^12^ further reinforces our observations and suggests a possible role for DGKζ in limiting T cell-dependent BM destruction.

The finding of CD69^+^ T cells in the BM of DGKζ^-/-^ mice at a very young age, together with the presence of Th1 and cytotoxic cytokines, suggest early presence of activated T cells. CD69 is an activation marker that is induced rapidly after antigen recognition and decays very rapidly in peripheral cells. In activated CD4 T cells, CD69 expression attenuates S1P1 expression and promotes homing into the BM, where CD4 T cells evolve into memory subsets.^47^ As a result, the BM is recognized as a central niche for long-term homing of memory T cells.^48^ The ability of CD69 to favor BM homing for CD4 T cells and attenuate S1P1 expression coincides with the greater CD4 T cell retention observed in the BM of DGKζ^-/-^ mice. DGKζ deficiency correlates with enhanced CD69 expression in CD4 and CD8 T cells.^44^ The reduction in naive populations and concomitant increase in memory T cells in PBL from DGKζ^-/-^ mice confirms self-activation, in accordance with the direct communication between the BM and peripheral blood.

Immune-dependent BM destruction in AA is attributed both to the direct and indirect effects of Th1 and CTL on HSC and other hematopoietic lineages.^49^ In healthy conditions, elevation of inflammatory cytokines such as IFNγ induce rapid differentiation of HSC to the myeloid lineage to respond to infections. Abnormal, chronic IFNγ elevation in AA impairs HSC continuous self-renewal leading to loss of stem precursors.^27^ The analysis of stem cell populations in DGKζ^-/-^ mice reveals an abnormal, high elevation of IFNγ concomitant with a significant decrease in all precursor populations and myeloid mature cells. In agreement, DGKζ^-/-^ mice have reduced numbers of circulating red cells, platelets, monocytes, and granulocytes. In addition to the loss of self-renewal abilities of the stem subsets, immune-dependent BM destruction is also linked to the direct induction of apoptosis. The increased apoptosis of erythroblasts observed even at a very early age, together with the lowered expression of cell cycle inhibitors probably linked to defective TPHO functions indicates that these are not opposite alternatives. These data coincides with the analysis of CD34^+^ cells from severe pediatric AA patients, which shows higher expression of apoptotic-related genes and an enhanced IFNγ signature.^50^ DGKζ^-/-^ mice show partial BM aplasia and diminished circulating platelets and granulocytes, but not severe anemia. It is plausible that low, chronic T cell activation leads to T cell exhaustion in a manner similar to the exhausted hyporesponsive T cells found in the BM of IFNγ-induced murine models of AA.^51^

The phenotype of DGKζ^-/-^ mice suggests BM destruction by autoreactive T cells. In agreement, infusion of DGKζ^-/-^ T splenocytes in histocompatibility-mismatched hosts accelerated BM damage when compared with splenocytes from WT mice. Cytokine analysis showed enhanced TNFα and IL-6 production in hosts infused with DGKζ^-/-^ cells. In GVHD models, transient IFNγ production associates with the initial highly proliferative phase whereas TNFα production correlates with the later more destructive phase of the disease.^42^ Our results suggest a temporality in the induction of BM failure as the result of a GVHD-like response; they suggest that DGKζ loss accelerates the onset of the reaction from the initial proliferative phase to the final destructive phase. This hypothesis coincides with recent studies in murine models of acute AA showing that TNFα, which is generated by host macrophages in response to infused T cells, accelerates immune-mediated BM destruction.^53^ Reduced IL-2 expression observed in mice that receive DGKζ-deficient cells is consistent with rapid onset of activation, leading to functional decline of antigen-specific effector T cells that could result in their depletion. Exhaustion of rapidly expanding effector T cell populations is said to explain stronger antitumor effects in xenotransplanted DGKζ^-/-^ mice.^6^ Little or no IFNγ production by BM T cells is also observed in very severe cases of AA, and is predictive of a lack of response to immunosuppressive therapy.^21^ Progressive T cell exhaustion would coincide with the lack of high IL-2 and TNFα expression observed in mature mice. It would also conform to reduced expression of IL-6, a pro-inflammatory cytokine that, as described for IFNγ and TNFα, regulates HSC differentiation.^52^

Despite intensive studies, understanding of the autoimmune mechanism in AA remains limited. Pesticides, toxic chemicals, and viral infections are hypothesized to confer higher risk, but acquired AA is largely considered idiopathic and the autoantigen remains unknown. We found no sex differences in the phenotype of DGKζ^-/-^ mice, whereas enhanced BM failure in older mice suggests cumulative damage. A progressive loss of BM function would agree with recent studies that cite normal platelet counts in DGKζ^-/-^ mice around 10 weeks of age.^54^ In contrast observations in older animals, our analysis confirms normal platelet counts between 10-15 weeks (not shown). The reported abnormal platelet responses^54^ could nonetheless be a result of defective differentiation. In this regard, all components of the trimeric GPIB/GPIX/GPV complex as well as GPVI, the collagen receptor on platelets, appear markedly downregulated in CD34^+^ gene profiling from pediatric severe AA patients.^50^

In summary, our study demonstrates that DGKζ protects the BM from autoimmune-mediated destruction, and suggests that DGKζ^-/-^ mice are a useful model to further explore the mechanisms that trigger this disease. Detection of DGKζ expression could also be an important biomarker of the T cell function in AA patients. Finally, a better understanding of the mechanism by which DGKζ protects the BM from autoimmune destruction could help to avoid the risks of aplastic anemia induced by treatment with immune checkpoint inhibitors.^55-57^

## Acknowledgments

We thank Cathy Mark for excellent editorial assistance and IM group members for helpful discussion. EA holds a predoctoral fellowship from the Spanish Ministry of Education. This work was supported in part by grants from the Aplastic Anemia and MDS International Foundation (AAMDSIF OPE01644), Spanish Ministry Economy and Competitiveness cofinanced by the European Regional Development Fund (BFU2016-77207-R) and the Madrid regional government (IMMUNOTHERCAM Consortium S2010/BMD-2326) to IM.

## Authorship contributions

IM, EA and MMS designed the research and wrote the manuscript; EA, MMS, MLS and RL performed the experiments; MMS and EA analyzed the results; MMS and EA designed the figures.

## Disclosure of conflicts of interest

The authors have no conflict of interests.

## Notes

### Competing Interest Statement

The authors have declared no competing interest.

