## Supplementary figures and images for "Diacylglycerol kinase ζ deficiency triggers early signs of aplastic anemia in mice"

### Supplemental Fig 1

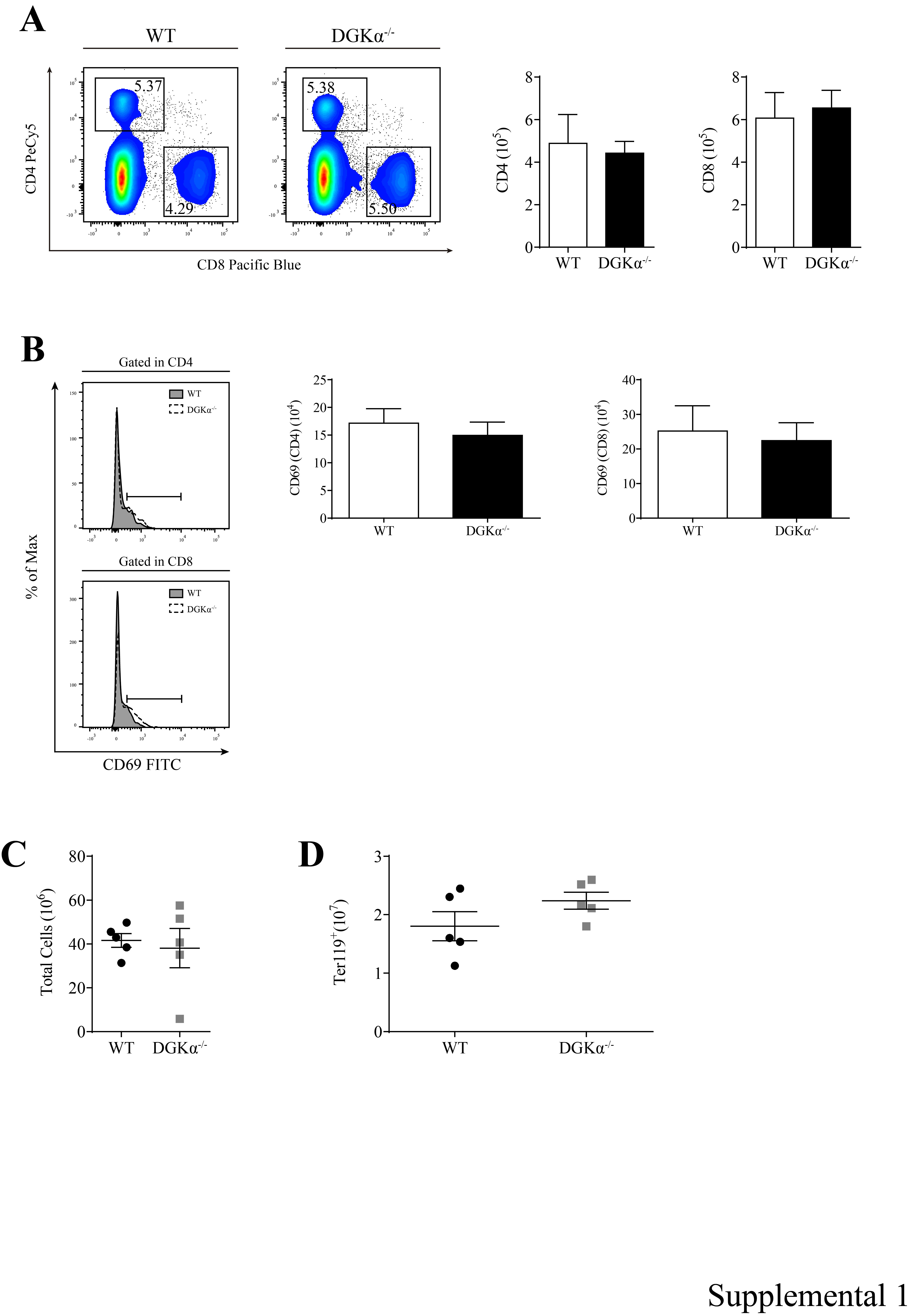
